# MicroRNA-200c suppresses epithelial-mesenchymal transition of ovarian cancer by targeting cofilin-2

**DOI:** 10.1101/449587

**Authors:** Xuechen Yu, Yuanzhen Zhang, Wei Zhang, Huijun Chen

## Abstract

This study investigated the effects of microRNA-200c (miR-200c) and cofilin-2 (CFL2) in regulating epithelial-mesenchymal transition (EMT) in ovarian cancer. The level of miR-200c was lower in invasive SKOV3 cells than that in non-invasive OVCAR3 cells, whereas CFL2 showed the opposite trend. Bioinformatics analysis and dual-luciferase reporter gene assays indicated that CFL2 was a direct target of miR-200c. Furthermore, SKOV3 and OVCAR3 cells were transfected with miR-200c mimic or inhibitor, pCDH-CFL2 (CFL2 overexpression), or CFL2 shRNA (CFL2 silencing). MiR-200c inhibition and CFL2 overexpression resulted in elevated levels of both CFL2 and vimentin while reducing E-cadherin expression. They also increased ovarian cancer cell invasion and migration *in vitro* and *in vivo* and increased the tumor volumes. Conversely, miR-200c mimic and CFL2 shRNA exerted the opposite effects as those aforementioned. In addition, the effects of pCDH-CFL2 and CFL2 shRNA were reversed by the miR-200c mimic and inhibitor, respectively. This finding suggested that miR-200c could be a potential tumor suppressor by targeting CFL2 in the EMT process.

## Introduction

Ovarian cancer is one of most common genital tract malignancy in females worldwide, ranking among the top 10 cancers in incidence and mortality globally (Sung et al., 2014). Ovarian cancer affects women when they are still young and has devastating effects with high personal, social, and economic cost (Bruggmann et al., 2017). The International Agency for Research on Cancer predicted that in 2012, 528,000 women were diagnosed with ovarian cancer, resulting in an estimated 266,000 deaths (Urban et al., 2016). Many other factors contribute to the progression of ovarian cancer. The process involves the transformation of normal ovarian epithelium into a ovarian intraepithelial pre-neoplasm, which is subsequently transformed into ovarian cancer. Clinical surgery and chemotherapy are not ideal for ovarian cancer patients because of the uncertainty in tumor characteristics. In addition, the mechanisms of tumor cell invasion and migration remain unclear.

Epithelial-mesenchymal transition (EMT) is a phenomenon wherein epithelial cells gain a mesenchymal phenotype (Chang et al., 2015). It is a potential mechanism by which cancer cells depart from the primary tumor and migrate to surrounding tissues and distant organs, resulting in enhanced invasion and metastasis (Fang et al., 2017). Concomitantly, EMT exhibits attenuated levels of epithelial markers such as E-cadherin and elevated levels of mesenchymal markers such as vimentin (Thiery et al., 2009). The control of the EMT process has been proposed as an approach against tumor invasion and metastasis.

MicroRNAs (miRNAs) are small (about 18-25 nucleotides) non-coding RNAs that control gene expression by binding with complementary sites in the 3’-untranslated regions (UTRs) of mRNA (Bartel, 2009; Moreno-Moya et al., 2014). Because of their abundance, mounting evidence has suggested that miRNAs play pivotal roles in a wide spectrum of biological processes, including EMT (Kim et al., 2011; Sekhon et al., 2016; Talbot et al., 2012; Yamada et al., 2013). MiR-200c is a member of the miR-200 family and is a marker of EMT in female reproductive cancers (Cochrane et al., 2010). Previous studies have shown that miR-200c regulates cell motility and invasiveness in breast, endometrial, and ovarian cancer cell lines (Ibrahim et al., 2015). Cofilin-2 (CFL2) is a small actin-binding protein expressed in skeletal and cardiac muscles that participates in regulating the actin cytoskeleton in cancer cell migration and invasion (Collazo et al., 2014; Yamaguchi and Condeelis, 2007). This study investigated whether miR-200c and CFL2 collectively regulate the EMT process and explored their involvement in the molecular mechanisms of ovarian cancer.

## Results

### Expression of miR-200c and CFL2 mRNA in invasive SKOV3 and non-invasive OVCAR3 cells

The mRNA level of miR-200c was markedly decreased in invasive SKOV3 cells compared to that in OVCAR3 cells (Figure 1A). Conversely, the mRNA level of CFL2 was increased in non-invasive SKOV3 cells compared to that in OVCAR3 cells (Figure 1B). These data suggested that the expression of CFL2 was higher in the invasive cell line and lower in the non-invasive cell line, whereas miR-200c showed the opposite trend.

**Figure 1.**
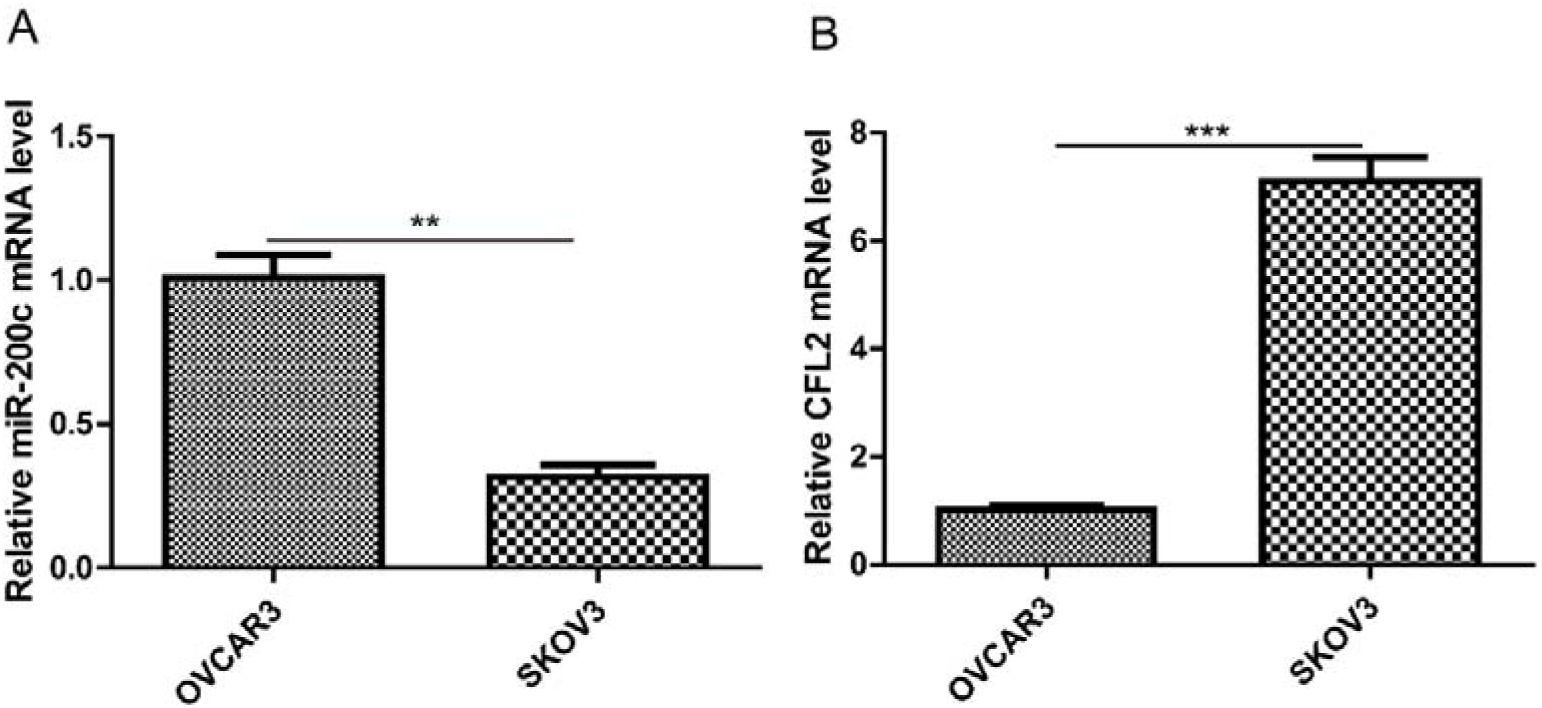
qRT-PCR assessment of miR-200c and CFL2 expression in OVCAR3 and SKOV3 cells. (A) miR-200c expression was significantly lower in SKOV3 cells relative to that in OVCAR3 cells. (B) CFL2 expression was significantly higher in SKOV3 cells relative to that in OVCAR3 cells. Each experiment was repeated three times, \*\*\**P* < 0.0001 and \*\**P* < 0.01.

### MiR-200c binds to 3’-UTR of CFL2

The miR-200c binding site in the CFL2 gene was located in the 3’-UTR, as revealed by the TargetScan software (Figure 2A). We verified these findings via dual-luciferase reporter gene assay (Figure 2B). Compared with the control group, the luciferase activity of the CFL2 3’-UTR-wt was suppressed in the miR-200c mimics group, whereas that of the CFL2 3’-UTR-mut showed no significant difference. This suggested that miR-200c specifically bound to CFL2 3’-UTR and negatively regulated CFL2 expression at the post-transcriptional level.

**Figure 2.**
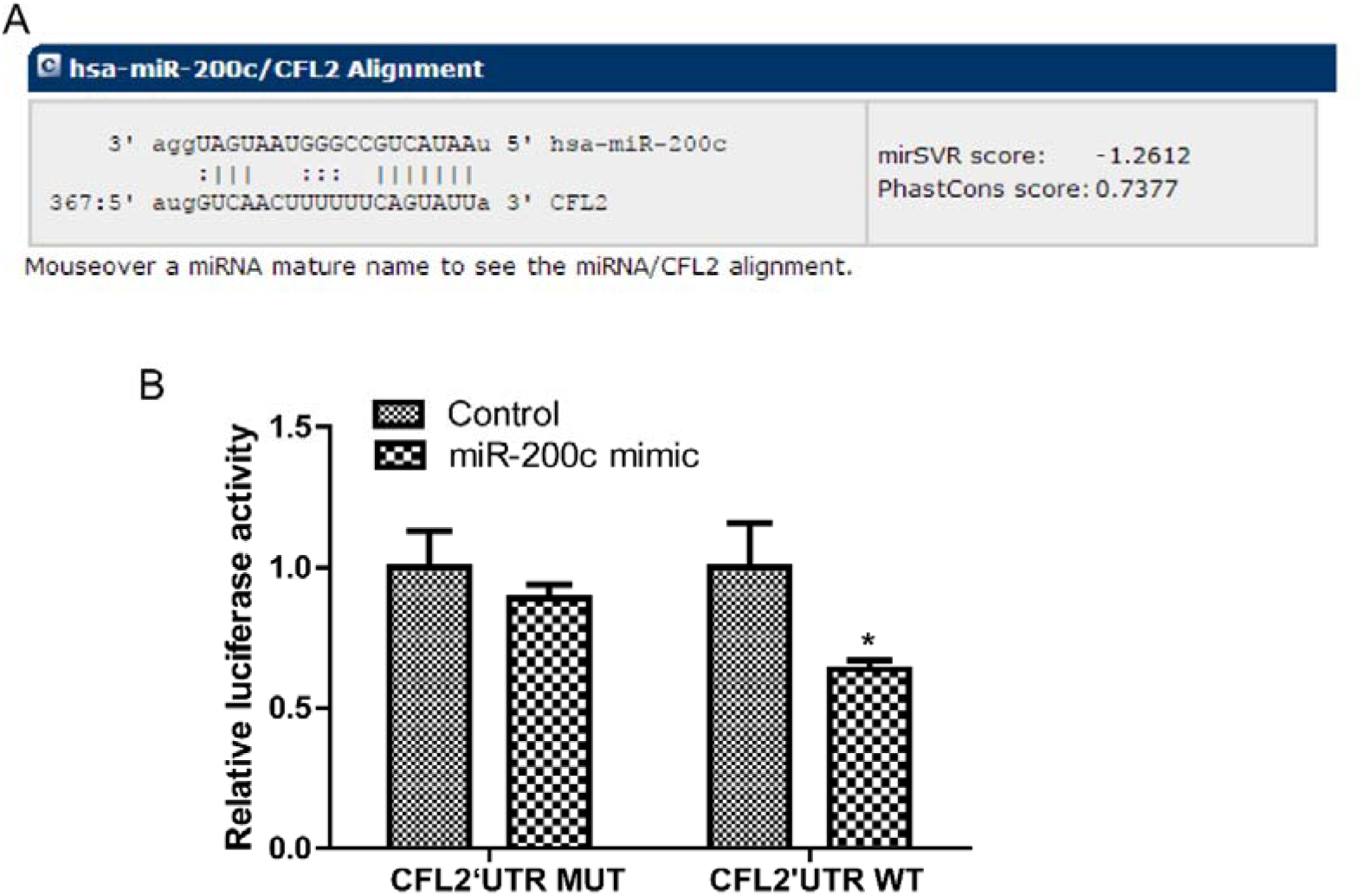
MiR-200c directly targets CFL2 by interacting with its 3’-UTR. (A) TargetScan prediction of miR-200c binding site in 3’-UTR of *CFL2* gene. (B) Dual-luciferase reporter gene assay showing miR-200c activity in the presence of CFL2 gene with wt and mut miR-200c binding site. Each experiment was repeated three times, \**P* < 0.05 compared with control.

### MiR-200c and CFL2 interaction regulated cell invasion and migration

The above findings indicated that a negative feedback was regulated between miR-200c and CFL2. To confirm this hypothesis, we conducted transwell assays to detect the invasion ability of SKOV3 and OVCAR3 cells with different transfections. The assay indicated that pCDH-CFL2 remarkably enhanced the invasiveness of non-invasive OVCAR3 cells, whereas cells treated with pCDH-CFL2 + miR-200c mimic exhibited a slight decrease in invasiveness compared with that of the pCDH-CFL2-treated cells (Figure 3). In contrast, miR-200c mimic markedly repressed the invasion of SKO3 cells, and CFL2 shRNA had the same effect (Figure 3). On the other hand, pCDH-CFL2-transfected OVCAR3 cells exhibited significantly increased migration ability, whereas significantly decreased migration was observed in OVCAR3 cells transfected with miR-200c mimic (Figure 4).

**Figure 3.**
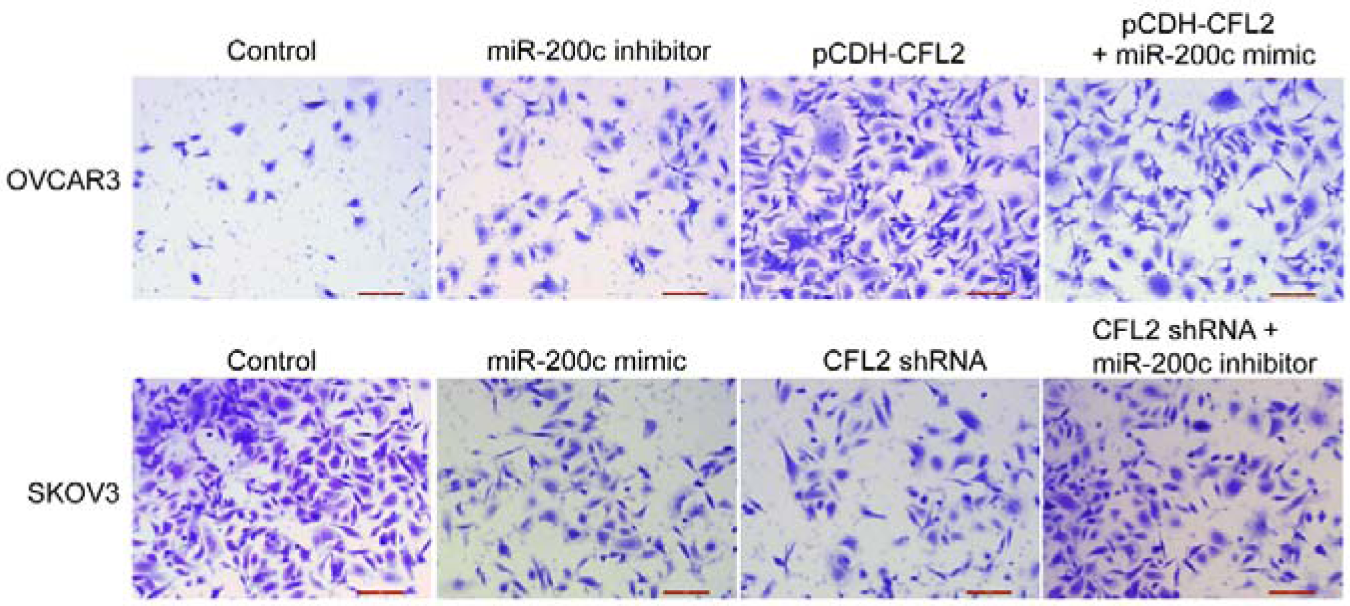
MiR-200c inhibited invasion and CFL2 overexpression promoted invasion of OVCAR3 and SKOV3 cells. Analysis of invasion of OVCAR3 and SKOV3 cells subjected to different transfections. Scale bar = 100 μm, magnification of 200×.

**Figure 4.**
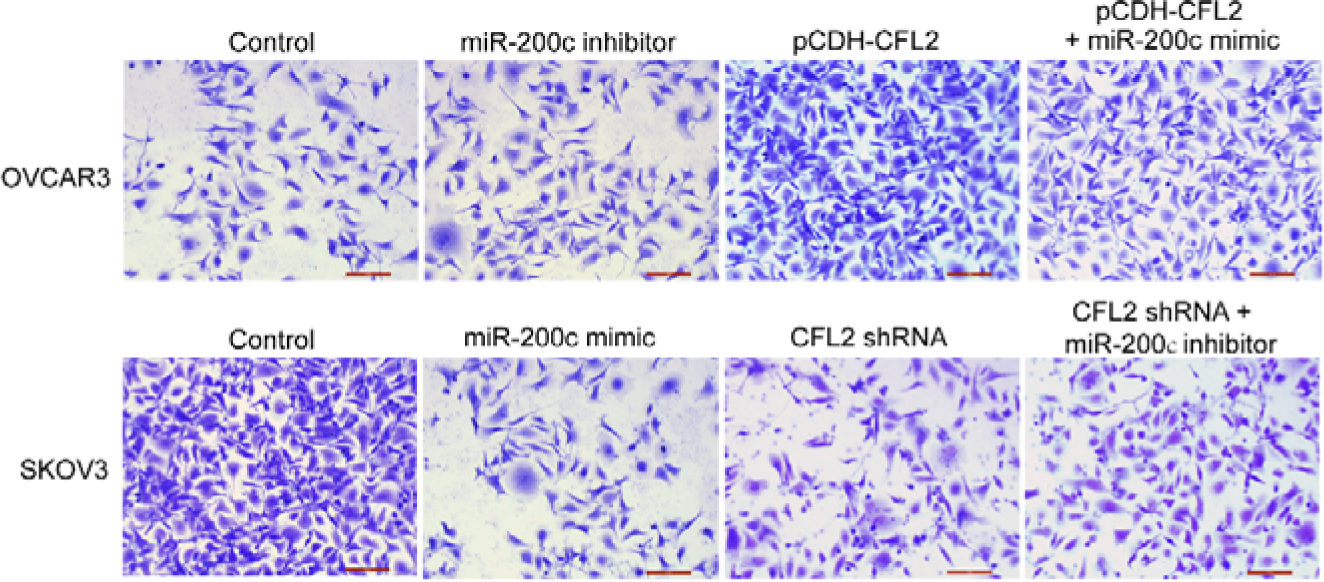
MiR-200c inhibited migration and CFL2 overexpression promoted migration of OVCAR3 and SKOV3 cells. Analysis of migration of OVCAR3 and SKOV3 cells subjected to different transfections. Scale bar = 100 μm, magnification of 200×.

### MiR-200c and CFL2 regulate vimentin and E-cadherin in ovarian cancer cells

Vimentin and E-cadherin are important proteins in relation to cell invasion and migration. In ovarian cell lines, the miR-200c inhibitor significantly increased the expression of CFL2 compared with that in control cells, and overexpression of the miR-200c mimic suppressed CFL2 expression. pCDH-CFL2 and CFL2 shRNA played essential roles in ovarian cancer cells by increasing or decreasing the expression of CFL2, respectively (Figure 5A). In pCDH-CFL2-treated OVCAR3 cells, vimentin was up-regulated and E-cadherin was down-regulated compared to the control. Comparatively, vimentin was down-regulated and E-cadherin was up-regulated when OVCAR3 cells were treated with miR-200c inhibitor (Figure 5B–D). In addition, OVCAR3 cells treated with pCDH-CFL2 + miR-200c mimic showed altered expressions of vimentin and E-cadherin compared with those in the pCDH-CFL2 group. On the other hand, vimentin was down-regulated and E-cadherin was up-regulated after the transfection of SKOV3 cells with either miR-200c mimic or CFL2 shRNA (Figure 5E–G). Meanwhile, SKOV3 cells transfected with CFL2 shRNA + miR-200c inhibitor showed altered expressions of vimentin and E-cadherin.

**Figure 5.**
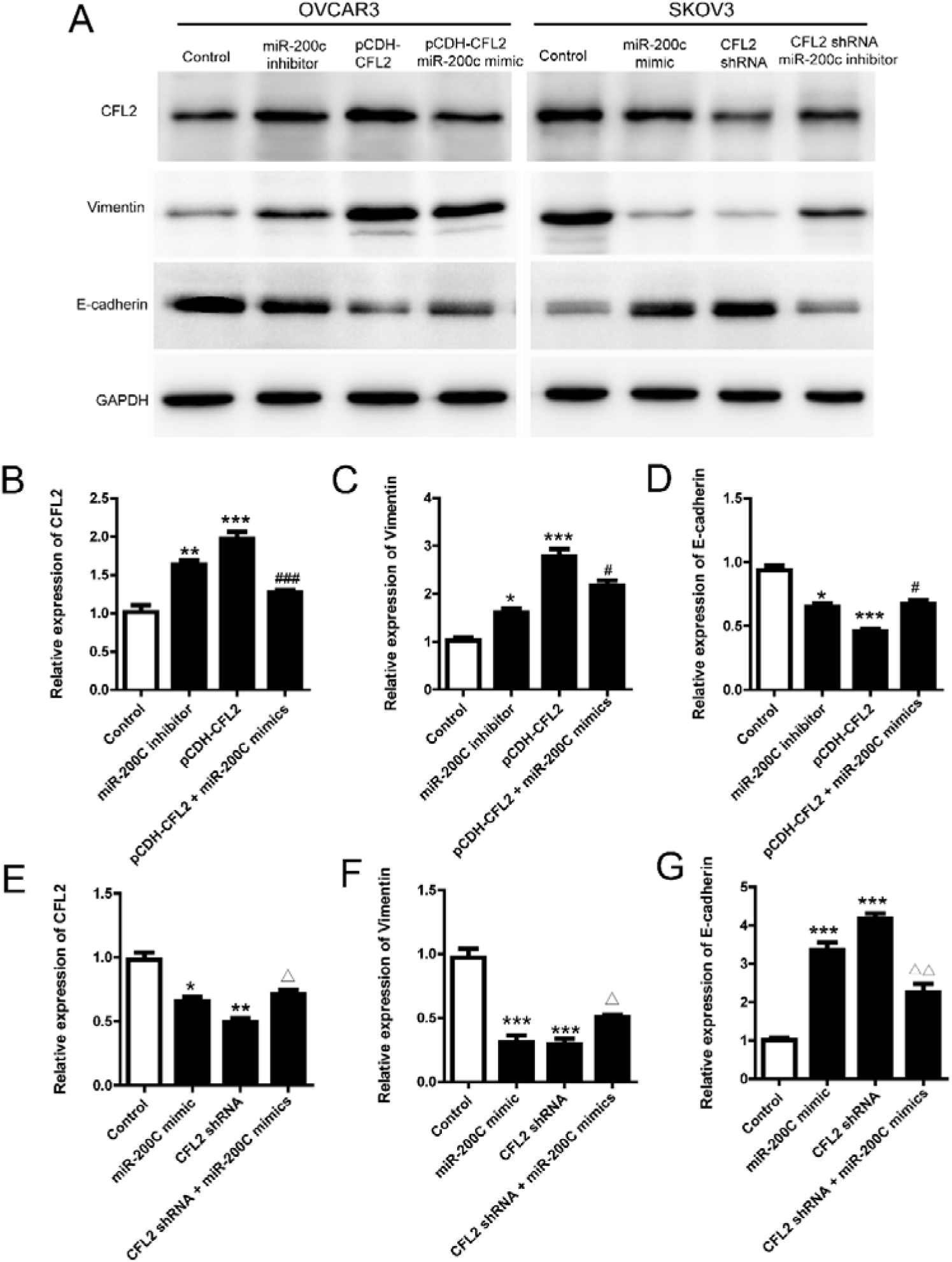
Western blot of EMT-related proteins.

(A) Expression of CFL2, vimentin, E-cadherin, and GAPDH in OVCAR3 and SKOV3 cells subjected to different transfections. Quantitative analysis of CFL2, vimentin, and E-cadherin expression in (B–D) OVCAR3 and (E–G) SKOV3 cells. GAPDH was used as a loading control. Each experiment was repeated three times, \*\*\**P* < 0.0001, \*\**P* < 0.01, and \**P* < 0.05 compared with control group; ^###^*P* < 0.0001 and ^#^*P* < 0.05 compared with pCDH-CFL2 group; ^ΔΔ^*P* < 0.01 and ^Δ^*P* < 0.05 compared with pCDH-CFL2 group.

### MiR-200c mimic significantly suppressed tumor growth in ovarian cancer xenografts in nude mice

To explore the antitumor efficacy of miR-200c in ovarian cancer cells, SKOV3 and OVCAR3 xenografts were established by injecting SKOV3 and OVCAR3 cells with different transfections. We found that CFL2 overexpression significantly promoted tumor growth in SKOV3-xenografted mice while the combination of CFL2 overexpression and miR-200c mimic slightly reduced tumor growth compared with that in the pCDH-CFL2 group (Figure 6A). Similarly, tumor growth in OVCAR3-xenografted mice was significantly inhibited in the miR-200c mimic and CFL2 shRNA groups compared with that in the control. The in vivo experiment suggested that the interaction between miR-200c and CLF2 regulated ovarian tumor growth.

**Figure 6.**
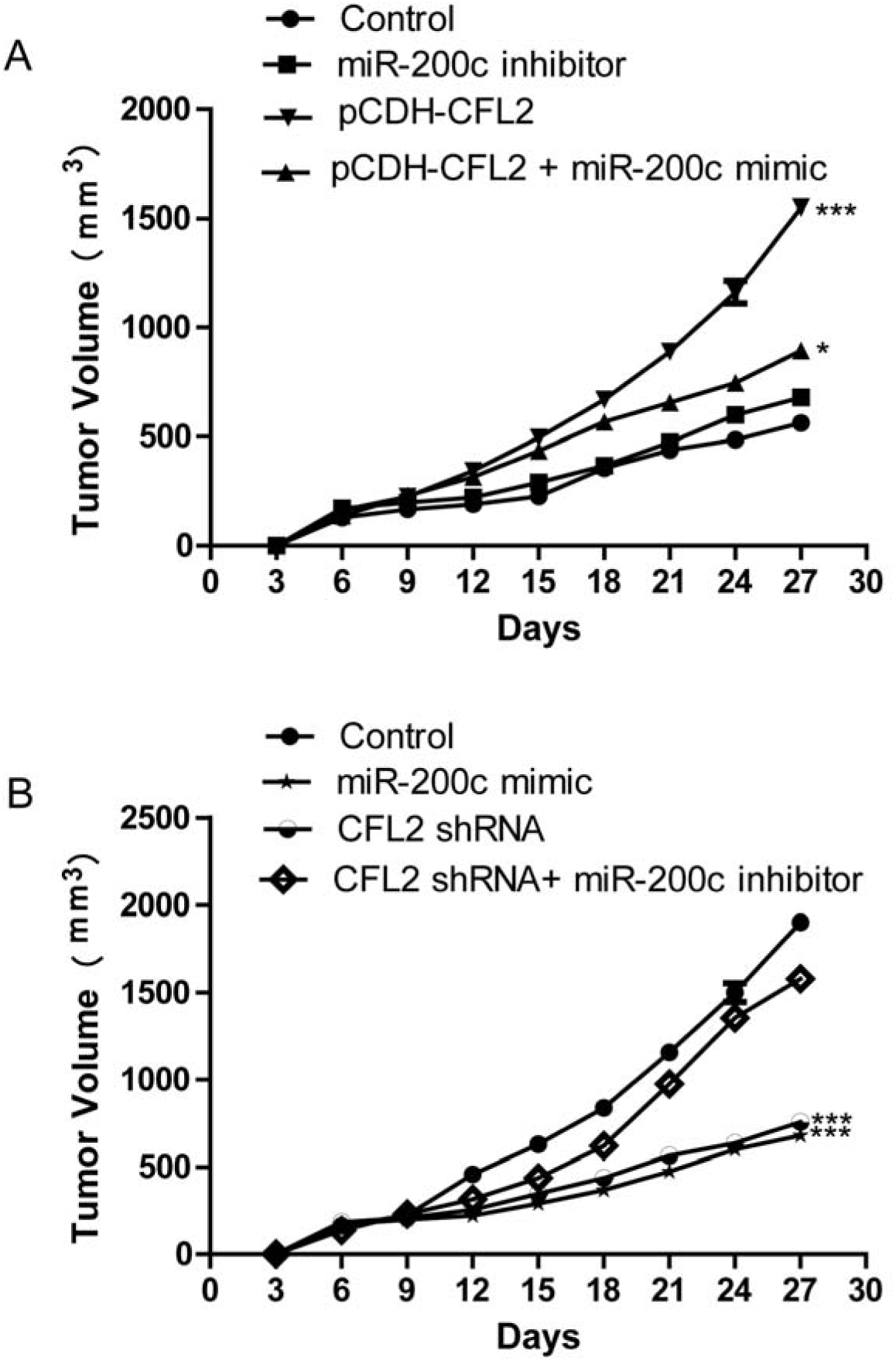
MiR-200c inhibited tumor growth in xenografted nude mice. (A) Total tumor volume in nude mice xenografts of control OVCAR3 cells and cells transfected with miR-200c inhibitor, pCDH-CFL2, or pCDH-CFL2 + miR-200c mimic. (B) Total tumor volume in nude mice xenografts of control SKOV3 cells and cells transfected with miR-200c mimic, CFL2-shRNA, or CFL2-shRNA + miR-200c inhibitor. \*\*\**P* < 0.0001 and \**P* < 0.05 compared with control group.

### MiR-200c and CFL2 regulated the expression of vimentin and E-cadherin in xenografts

To further investigate the role of EMT-relevant markers in ovarian cancer, immunohistochemistry was employed to detect vimentin and E-cadherin expression in tumor tissues extracted from nude mice. The results showed that the expressions of vimentin and CFL2 in the pCDH-CFL2 treatment group were obviously higher than those in the control and miR-200c inhibitor groups. However, E-cadherin expression in the pCDH-CFL2 group was significantly lower than that in the control and miR-200c inhibitor groups in OVCAR3-xenografted tumor tissues (Figure 7A). On the other hand, miR-200c mimic and CFL2 shRNA promoted the expression of E-cadherin and suppressed vimentin and CFL2 expression compared with those in the control group in SKOV3-xenografted tumor tissues (Figure 7B). These findings suggest that the effective inhibition of ovarian cancer growth by miR-200c may be partly achieved by regulating the expression of vimentin and E-cadherin.

**Figure 7.**
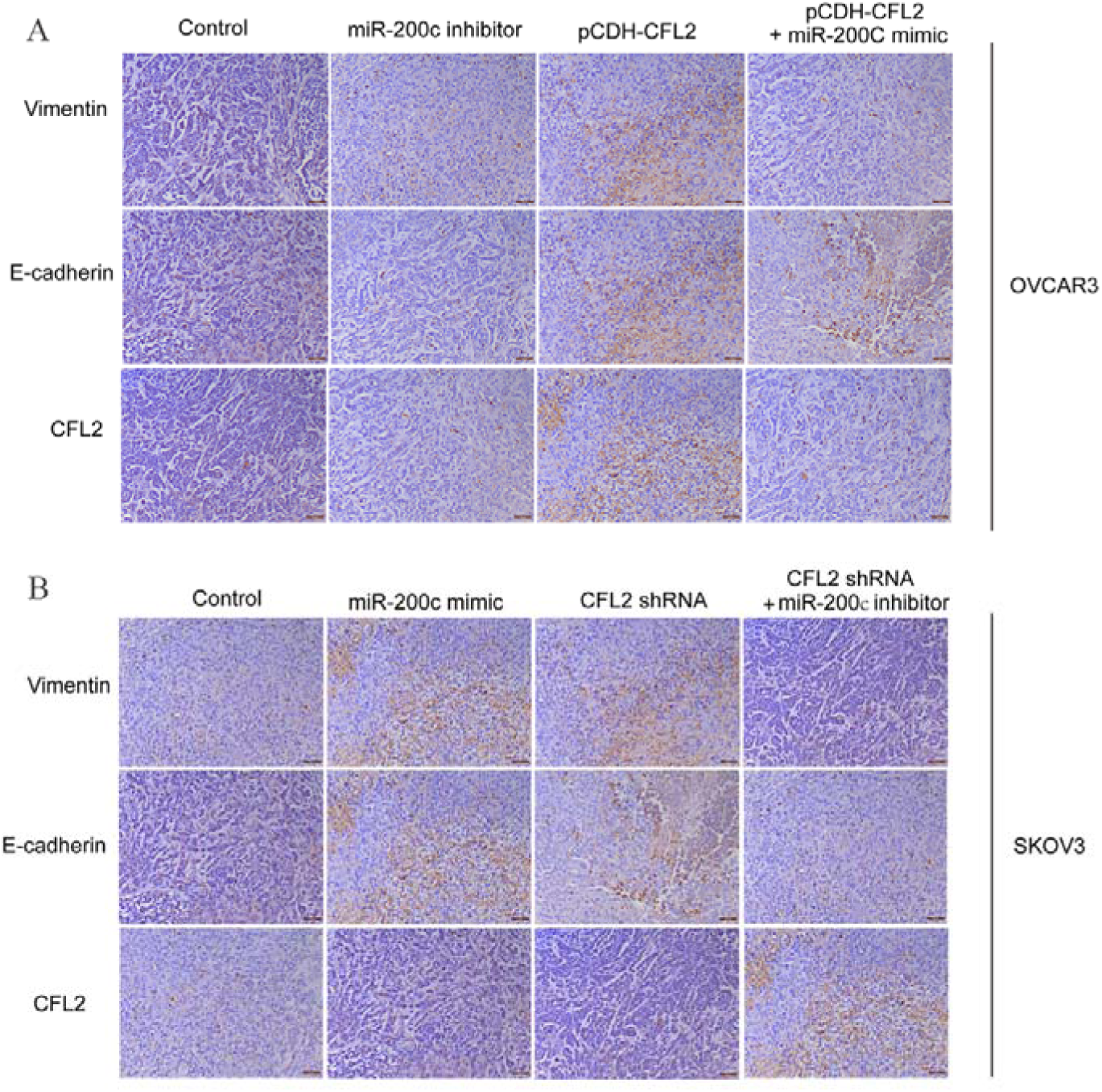
Immunohistochemistry of tumor tissues from nude mice subjected to different transfections. (A) Expression of vimentin, E-cadherin, and CFL2 in tumor tissue of control OVCAR3 cells and cells transfected with miR-200c inhibitor, pCDH-CFL2, or pCDH-CFL2 + miR-200c mimic in nude mice. (B) Expression of vimentin, E-cadherin, and CFL2 in tumor tissue of control SKOV3 cells and cells transfected with miR-200c mimic, CFL2 shRNA, or CFL2 shRNA + miR-200c inhibitor in nude mice. Scale bar = 100 μm.

## Discussion

Currently, the molecular mechanisms through which miRNA mediates EMT in ovarian cancer is not yet fully elucidated. A full understanding of these mechanisms is necessary to improve ovarian cancer treatment. Increasing evidence has suggested that the expression of miRNA is frequently associated with EMT (Braun et al., 2010; Zheng et al., 2016). In our study of the regulation and expression of miR-200c, CFL2 was revealed as a new target gene regulated by miR-200c. In particular, miR-200c-mediated down-regulation of CFL2 played a crucial role in the reduction of EMT in ovarian cancer cells and the inhibition of tumor cell growth in vitro and in vivo. The present study demonstrated that miR-200c and CFL2 regulated the invasiveness and migration of ovarian cancer cells. We observed that the non-invasive OVCAR3 cells had high expression of miR-200c and low expression of CFL2, whereas the invasive SKOV3 cells showed the opposite trend. This finding suggested that miR-200c suppressed the invasiveness of ovarian cancer cells, which was consistent with the results of a previous report (Sulaiman et al., 2016).

It has been reported that miR-21 treatment of cholangiocarcinoma xenografts resulted in elevated levels of vimentin and reduced levels of E-cadherin (Liu et al., 2015). In our study, miR-200c mimics or inhibitors were transfected into OVCAR3 and SKOV3 cells. The EMT phenotype of reduced E-cadherin and elevated vimentin was induced by miR-200c inhibitor treatment and reversed by miR-200c mimic. In addition, CFL2 was silenced or overexpressed using CFL2 shRNA or pCDH-CFL2, respectively, in OVCAR3 and SKOV3 cells. Furthermore, E-cadherin and vimentin expression showed opposite trends with CFL2 shRNA or pCDH-CFL2 treatment. Liu et al. reported that miR-200c mimic significantly reduced EMT, invasion, and migration while the inhibitor generated the opposite effects (Liu et al., 2017), which were consistent with our in vitro and in vivo experimental results. Furthermore, Zhang et al. suggested that miR-200c inhibited the proliferation, migration, and invasion of gastric cancer cells (Zhang et al., 2017).

Our study was not only based on cell lines but also involved tumor-bearing nude mice. We found that the expression of miR-200c was high in non-invasive ovarian cancer cells and low in invasive ovarian cancer cells, and CFL2 exhibited the opposite trend. MiR-200c mimic inhibited the invasion and migration of SKOV3 and OVCAR3 cells, suggesting that miR-200c acts as a tumor suppressor. CFL2 was identified as a target of miR-200c and was down-regulated by miR-200c. It was involved in the miR-200c-mediated regulation of ovarian cancer cell invasion and migration in vitro and in vivo. High CFL2 expression promoted cell invasion and migration, and miR-200c mimic reduced the effect of CFL2 overexpression. Similarly, miR-200c inhibitor suppressed the effect of CFL2 shRNA on the inhibition of cell invasion and migration, suggesting that the reduction of miR-200c attenuated the inhibitory effect of CFL2 shRNA blockade on the properties of ovarian cancer cells.

In conclusion, our study identified CFL2 as a new target of miR-200c in regulating ovarian cancer cell invasion and migration in vitro and in vivo, potentially laying the foundation for the clinical application of CFL2.

## Materials and Methods

### Cell culture

Human ovarian cancer cell lines OVCAR3 and SKOV3 were purchased from the American Type Culture Collection and cultured in RPMI-1640 medium supplemented with 10 % fetal bovine serum (FBS, Corning) and 100 μg/mL penicillin/streptomycin (Sigma). All cell were maintained at 37 °C in a humidified atmosphere composed of 5 % CO_2_ and 95 % air. Cells in the logarithmic growth phase were used for subsequent research.

### Plasmids

pSIpCDH-CMV-MCS-EF1-CopGFP-T2A-Puro (pCDH-CFL2, containing CFL2 sequence) and pCMV-GFP-shCFL2 (short hairpin CFL2, containing shCFL2 sequence) plasmids were sequenced in both forward and reverse directions by RiboBio (Guangzhou, China).

### Cell transfection and grouping

SKOV3 cells were transfected with miR-200c mimic (synthetic miR-200c analogs purchased from RiboBio), pCMV-GFP-shCFL2 (CFL2 silencing), or pCMV-GFP-shCFL2+miR-200c inhibitor (RiboBio, Guangzhou), and non-transfected cells were used as a control. OVCAR3 cells were transfected with miR-200c inhibitor, pCDH-CFL2 (CFL2 overexpression), or pCDH-CFL2+miR-200c mimic, and non-transfected cells were used as a control. Before transfection, an appropriate number of cells were inoculated in two 6-well culture plates with RPMI-1640 complete medium to reach a cell density of 30–50 % until transfection. The cells were mixed with 100 μL of serum-free culture medium in eppendorf tubes and incubated with 5 µL of Lipofectamine 2000 (Invitrogen, New York, USA) for 5 min at room temperature. Subsequently, the mimics, inhibitors, and plasmids were added to each group at a final concentration of 500 μL/well and mixed gently. The plates were incubated in an incubator containing 5% CO_2_ at 37 °C for 6 h. Thereafter, the old medium was replaced by complete medium containing 10 % FBS for continuous culture. The cells were collected 48 h after transfection and subjected to western blot and quantitative reverse-transcription polymerase chain reaction (qRT-PCR) for migration and invasion analysis.

### Dual-luciferase report analysis

Bioinformatics was performed to predict the close relationship between miR-200c and CFL2. The miRNA-200c binding site on the 3’-UTR of CFL2 mRNA was predicted using TargetScan (www.targetscan.org). To confirm miR-200c binding to the 3’-UTR site in the CFL2 gene, genomic DNA was extracted from SKOV3 and OVCAR3 cells. The CFL2 3’-UTR-wildtype (wt) was bound to the miRNA-200c-wt binding site (5’-CTAGCTCCATTTCTCCAGCTCAGTCCATTGGAATAGTATT AG GTTTTGGTTTTTTGTTGTATTTCCCCCT-3’) and the CFL2 3’-UTR-mutant (mut) was bound to the miRNA-200c-mut binding site (5’-CTAGCTCCATTTCTCCAGCTCAGTCCATTGGAATACTAATAGGTTTTGGTTTTTTGTTGTATTTCCCCCT-3’). Both CFL2 3’-UTR-wt and CFL2 3’-UTR-mut were cloned into pMir-GLO vectors (Promega Company, USA) with luciferase reporters and transfected into HEK-293T cells. The dual-luciferase reporter assay was performed to test the luciferase activity of the samples 48 h after transfection according to the manufacturer’s instructions (Promega Company, USA).

### Transwell migration and invasion assays

Transfected SKOV3 and OVCAR3 cells were harvested in the logarithmic growth phase by digesting with 0.25 % trypsin when they reached 90 % confluence. In vitro cell migration and invasion assays were performed using Transwell chambers with polyethylene terephthalate membranes (24-well inserts, 8.0 μm; Corning). For the migration assay, 3 × 10^4^ cells were added to the top chamber. For the invasion assay, approximately 5 × 10^4^ cells were seeded into the top chamber coated with Matrigel (BD Biosciences). Complete medium was added to the bottom wells to stimulate migration or invasion. After the cells were incubated for 48 h, they were stained with 0.1 % crystal violet. Five fields per filter were counted using Image-Pro Plus software.

### qRT-PCR

Transfected SKOV3 and OVCAR3 cells were harvested in the logarithmic growth phase by digesting with 0.25 % trypsin when they reached 90 % confluence. Then total RNA was extracted using Trizol reagent (Invitrogen, USA) according to the manufacturer’s protocol. Complementary DNA was reverse-transcribed using a reverse transcription kit (Invitrogen, USA) and qRT-PCR was performed using SYBR Green Supermix (Bio-Rad) on a CFX96™ Real Time PCR system (Bio-Rad). The conditions of qRT-PCR were as follows: 95 °C for 5 min, 94 °C for 30 s, 56 °C for 30 s, and 72 °C for 30 s, 35 cycles, to obtain fluorescence intensity. GAPDH was used as an internal control in all experiments. The sequences of PCR primers are listed as follows: miRNA-200c, 5’-GTCGTATCCAGTGCGTGTCGTGGAGTCGGCAATGCACTGGATACGACTCCATC-3’; E-cadherin, forward 5’-TCGACACCCGATTCAAA GTGG-3’ and reverse 5’-TTCCAGAAACGGAGGCCTGAT-3’; vimentin, forward 5’-CCTTGACATTGAGATTGCCA-3’ and reverse 5’-GTATCAACCAGAGGGAGT GA-3’; GAPDH, forward 5’-GCTGCCCAACGCACCGAATA-3’ and reverse 5’-GAG TCAACGGATTTGGT CGT-3’. Fold changes in gene expression were calculated by a comparative threshold cycle (Ct) method using the formula 2^-ΔΔCt^.

### Western blot

Transfected SKOV3 and OVCAR3 cells were harvested in the logarithmic growth phase by digesting with 0.25 % trypsin when they reached 90 % confluence. The cells were homogenized in a lysate buffer and then centrifuged at 4 °C for 25 min. Protein concentrations were measured and 60 µg of proteins were fractionated by 7.5 % or 10 % sodium dodecyl sulfate-polyacrylamide gel electrophoresis and transferred to polyvinylidene fluoride membranes (Millipore, Billerica, MA, USA). The membranes were blocked with 5 % skim milk, probed with primary antibodies overnight at 4 °C, and incubated with horseradish peroxidase-conjugated secondary antibody.

### Establishment of ovarian cancer-bearing nude mouse model

Mice were group-housed in polypropylene cages and maintained on a 12-h light/dark cycle at 22–25 °C and 50–65 % relative humidity. Transfected SKOV3 and OVCAR3 cells were harvested in the logarithmic growth phase by digesting with 0.25 % trypsin when they reached 90 % confluence. Cells (1 × 10^6^/200 μL) were inoculated in BALB/c nude mice subcutaneously. The maximum diameter (a) and maximum transverse diameter (b) of tumors were measured every three days and the tumor volume (V = ab^2^/2) was calculated. After 4 weeks, all mice were sacrificed by ovarian dislocation. The tumors were surgically removed and weighed, and the final tumor volumes were calculated. All animal experiments were approved by the Animal Experiment Ethics Committee and performed in accordance with the Regulations for Animal Experiments and Related Activities.

### Immunohistochemistry

Tumor tissues were collected after the mice were sacrificed and immunohistochemistry was performed according to a reported method. Briefly, 5-μm-thick serial tumor tissue sections (paraffin-embedded) were de-waxed in xylene and rehydrated in graded alcohol, after which endogenous peroxidase activity was quenched by incubating the sections in 3 % (v/v) H_2_O_2_ in methanol. Antigen retrieval was performed by incubating the sections in citrate buffer (pH 6.0). Non-specific binding was blocked by 5 % bovine serum albumin. After overnight incubation with the primary antibodies (E-cadherin and vimentin) at 4 °C, the sections were washed with phosphate-buffered saline (PBS) and incubated for 1 h with biotinylated goat anti-rabbit immunoglobulin G diluted at 1:200 in PBS. The slices were mounted with neutral gum for microscopic examination, and images were collected using a Nikon eclipse Ti-S microscope at 200× magnification. Cells with brown granules in the cytoplasm or nucleolus were considered positive.

### Statistical analysis

Statistical analysis was performed with GraphPad Prism 5 for Windows (GraphPad Software). Data are presented as the mean ± standard error of the mean. Numerical values represent the mean of three independent experiments. Statistical evaluation of the data was performed using Student’s t-test and two-way analysis of variance for multiple comparisons. Significant difference is defined at a P value of <0.05.

## Acknowledgement

This work is jointly supported by Fundamental Research Funds for the Central Universities No. 2042017kf0146.

**Abbreviations**
miR-200c: microRNA-200c
CFL2: cofilin-2
EMT: epithelial-mesenchymal transition
UTRs: untranslated regions
FBS: fetal bovine serum
qRT-PCR: quantitative reverse-transcription polymerase chain reaction
PBS: phosphate-buffered saline

